# *Sulfuriferula* spp. from sulfide mineral weathering environments have diverse sulfur- and iron-cycling capabilities

**DOI:** 10.1101/2025.01.29.635580

**Authors:** Kathryn K. Hobart, Gabriel M. Walker, Joshua M. Feinberg, Jake V. Bailey, Daniel S. Jones

## Abstract

Microorganisms are important catalysts for the oxidation of reduced inorganic sulfur compounds. One environmentally important source of reduced sulfur is metal sulfide minerals that occur in economic mineral deposits and mine waste. Previous research found that *Sulfuriferula* spp. were abundant and active in long-term weathering experiments with simulated waste rock and tailings from the Duluth Complex, Northern Minnesota. We therefore isolated several strains of *Sulfuriferula* spp. from these long-term experiments and characterized their metabolic and genomic properties to provide insight into microbe-mineral interactions and the microbial biogeochemistry in these and other moderately acidic to circumneutral environments. The *Sulfuriferula* strains are all obligate chemolithoautotrophs capable of oxidizing inorganic sulfur compounds and ferrous iron. The strains grew over different pH ranges, but all grew between pH 4.5-7, matching the weathering conditions of the Duluth Complex rocks. All strains grew on the iron-sulfide mineral pyrrhotite (Fe_1-x_S, 0 < x < 0.125) as the sole energy source, as well as hydrogen sulfide and thiosulfate, which are products of sulfide mineral breakdown. Despite their metabolic similarities, each strain encodes a distinct pathway for the oxidation of reduced inorganic sulfur compounds as well as differences in nitrogen metabolism that reveal diverse genomic capabilities among the group. Our results show that *Sulfuriferula* spp. are primary producers that likely play a role in sulfide mineral breakdown in moderately acidic to circumneutral mine waste, and the metabolic diversity within the genus likely explains their success in sulfide mineral-rich and other sulfidic environments.

**Importance:** Metal sulfide minerals such as pyrite and pyrrhotite are one of the main sources of reduced sulfur in the global sulfur cycle. The chemolithotrophic microorganisms that break down these minerals in natural and engineered settings are catalysts for biogeochemical sulfur cycling and have important applications in biotechnological processes such as biomining or bioremediation. *Sulfuriferula* is a recently described genus of sulfur oxidizing bacteria that are abundant primary producers in diverse terrestrial environments, including waste rock and tailings from metal mining operations. In this study, we explored the genomic and metabolic properties of new isolates from this genus, and the implications for their ecophysiology and biotechnological potential in ore and waste from economic mineral deposits.

## Introduction

The oxidation of reduced inorganic sulfur compounds is a crucial step in the global sulfur cycle. Sources of reduced sulfur include metal sulfide minerals that are present in sedimentary, igneous, and metamorphic rocks, dissolved and gaseous hydrogen sulfide species that are common in anoxic environments, and compounds with intermediate valences that include both short-lived compounds like thiosulfate (S_2_O_3_^2-^) and tetrathionate (S_4_O_6_^2-^) and more stable species like elemental sulfur. Reduced inorganic sulfur compounds are a dynamic source of energy for modern microorganisms that drive the oxidation of these compounds in diverse environmental conditions (1–6), and have been important throughout Earth’s history (7, 8). Many of these inorganic-sulfur-oxidizing bacteria and archaea are primary producers that fix carbon and nitrogen, and are abundant in environments with a range of pH conditions (from pH less than 1 to above 10) and temperatures (below 10 to over 90℃) (9). Terrestrial environments are particularly diverse settings for sulfur cycling because sulfur sources can be heterogeneously distributed. Steep gradients in environmental conditions like pH, temperature, and redox potential provide many distinct niches for different sulfur cycling organisms to thrive (1, 10–13).

*Sulfuriferula* are a recently described genus of sulfur-oxidizing bacteria. The first *Sulfuriferula* strains were isolated from a uranium mine using galena (PbS) as the sole electron donor, and was originally named ‘*Thiobacillus plumbophilus*’ (14). The genus *Sulfuriferula* was formally described in 2015 within the new family *Sulfuricellaceae*, which included members reclassified from the polyphyletic genus *Thiobacillus* (15). Members of the genus *Sulfuriferula* can grow autotrophically on inorganic sulfur compounds, and some can grow organoheterotrophically on organic substrates (15). Formally described members of the genus *Sulfuriferula* include *S. multivorans*, which was isolated from a freshwater lake in Hokkaido, Japan (15); *S. thiophila*, which was isolated from a storage tank at a public hot spring bath in Yamanashi Prefecture, Japan (16); *S. nivalis*, isolated from snow collected at the shore of a pond in Daisetsuzan National Park, Japan (17); and *S. plumbiphila*, the original “Thiobacillus plumbophilus” isolate that was formally described and later renamed (14–16). All species have been reported to grow aerobically on thiosulfate, tetrathionate, and elemental sulfur, while only *S. multivorans* can grow heterotrophically and anaerobically with nitrate as an electron acceptor (15) and *S. plumbiphila* can also grow autotrophically on H_2_ and PbS (14). *Sulfuriferula* are now known from diverse sulfur-rich terrestrial environments, and relatives of *Sulfuriferula* in metagenomic and 16S rRNA gene datasets have been found in metal- and sulfide-rich mine tailings and drainage (18–26), sewer and groundwater environments (27–29), hot springs and volcanic deposits (30–32), seawater-influenced sulfide caves (33), and low-temperature lake bottom sediments (18, 34, 35).

Here we describe new strains of *Sulfuriferula* that were isolated from weathered rocks from the sulfide mineral-bearing Duluth Complex in Northern Minnesota. The Duluth Complex contains copper, nickel, and platinum group element (Cu-Ni-PGE) deposits that may represent the world’s largest undeveloped economic Cu-Ni-PGE resource (36). The most abundant sulfide mineral in these rocks is pyrrhotite (Fe_1-x_S, 0 < x < 0.125), followed by chalcopyrite (CuFeS_2_), cubanite (CuFe_2_S_3_), and pentlandite ((Fe, Ni)_9_S_8_), with smaller amounts of other Cu, Ni, and PGE-bearing sulfide minerals (37). Sulfur- and iron-oxidizing microorganisms are known to accelerate the dissolution of sulfide minerals under highly acidic conditions (38). However, in contrast to many other well-studied ore and mine-waste environments where extreme acidity is generated, the relatively low sulfide mineral content (usually <1% total sulfur) and buffering capacity of the surrounding silicate minerals means that leachate from Duluth Complex waste rock and tailings is moderately acidic and rarely reaches pH values of less than 4 (39, 40). The role of microorganisms in this mildly acidic environment is not as well understood as in extremely acidic systems.

Based on rRNA gene and transcript sequencing, *Sulfuriferula* spp. were some of the most abundant and active organisms in experimentally weathered Duluth Complex rocks and tailings (18), and were also found (although at lower abundances) in naturally-weathered Duluth Complex outcrops (41), suggesting that they are important members of the rock weathering community and may also be valuable for developing biological solutions to managing mine waste and water from proposed mines in the region. To learn more about these important organisms and the role they play in the biogeochemistry of Duluth Complex mine waste and water, we isolated and characterized four strains of *Sulfuriferula* from experimentally weathered Duluth Complex materials. We describe the genomic and physiological properties of the strains and connect their diverse metabolic capabilities to their ecophysiology in the rock weathering environment.

## RESULTS

### Isolation and identification of *Sulfuriferula*

Strains of *Sulfuriferula* were isolated from weathering experiments on mine tailings and crushed rock from copper-nickel deposits in the Duluth Complex, northeastern Minnesota. Strains AH1, GW1, and GW6 were isolated from tailings and crushed rock had been experimentally weathered for more than 12 years using a laboratory humidity cell apparatus, and strain HF6a was isolated from an experimental field pile that had been weathering outside for more than 20 years (39, 40). Sixteen isolates were obtained, and four phylogenetically distinct strains, referred to as AH1, GW1, GW6, and HF6a, were selected for further study. Strains AH1, GW1, and HF6a were enriched and isolated on solid thiosulfate media that contained 6 mM ammonium, while GW6 was enriched and isolated on media with urea as the sole provided N source.

Phylogenetic analysis of 16S rRNA gene sequences from these isolates confirm that all cluster with described *Sulfuriferula* species (Figure 1). Strains GW1 and HF6a are most closely related to *S. multivorans* (15), strain GW6 is most closely related to *S. plumbiphila* (14, 15), and strain AH1 is located in its own clade located at the base of the *Sulfuriferula* clade near *S. thiophila* and *S. nivalis* (16, 17, 42). Strains had either two (GW1, GW6, and HF6a) or three (AH1) *rrn* operons with identical 16S rRNA genes (Figure 1).

**Figure 1.**
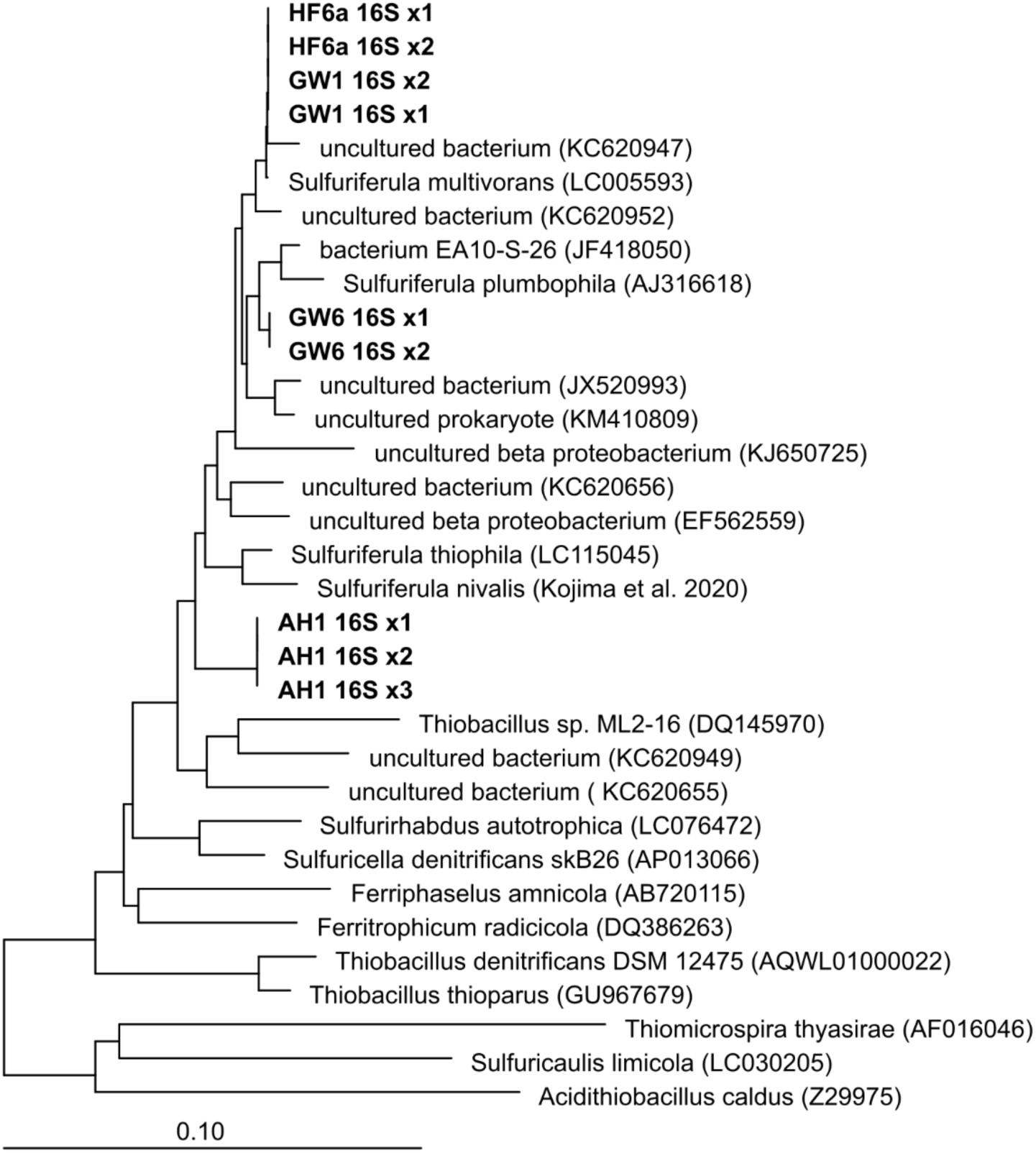
Neighbor-joining phylogenetic tree of 16S rRNA gene sequences, showing the relationship of the four Sulfuriferula isolates to other members of family Sulfuricellaceae. Names in bold font are sequences from this study, and the numbers after each name (x1, x2) indicate rRNA gene copies from multiple rrn operons in the genome.

### Growth characteristics

All four *Sulfuriferula* isolates grew on autotrophic media with thiosulfate (S_2_O_3_^2^-) as the only available electron donor. No growth was detected on tetrathionate (S_4_O_6_^2-^), sulfite (SO_3_^2-^), or elemental sulfur (S^0^) on the media used here. All strains also grew on pyrrhotite (Fe_1-x_S, 0 ≤ x ≤ 0.125) as the sole electron donor, and on dissolved sulfide (H_2_S(*aq*)) in gradient tubes. When grown on thiosulfate, strain AH1 accumulates sulfate and H^+^ ions in the growth media, consistent with complete thiosulfate oxidation to sulfate (Figure S1). In addition to reduced inorganic sulfur compounds, all strains grew on ferrous iron (Fe^2+^) in gradient tubes (Supplementary Figure S1). However, we did not observe anaerobic growth on either H_2_S(*aq*) or S_2_O_3_^2-^ with NO_3_^-^ as an electron acceptor under the conditions tested here, or when only hydrogen (H_2_) was provided as an electron donor under aerobic or anaerobic conditions.

Growth was not observed for any strains exclusively on LB broth or 1:10 diluted LB broth. No increase in growth was observed when 5mM glucose, acetate, or succinate was added to the thiosulfate media, suggesting that these strains are obligate autotrophs and are sensitive to the presence of organics. Consistent with this, growth rate decreased in media buffered with organic buffers rather than phosphate. Growth occurred in MES- and HEPES- but not citrate-buffered media. However, growth was enhanced by adding 0.1% w/v yeast extract was added to the growth media.

Each strain grew at slightly different pH ranges. Strain AH1 had the widest range, which grew between pH 4.5 and 8. HF6a was most acid tolerant than the other strains, growing between pH 4 and 7, while strains GW1 and GW6 were less so, growing between pH 5 and 8 and 6 and 8 respectively. During growth on thiosulfate, pH consistently initially increased and then decreased (e.g., Supplementary Figure S2).

### Whole genome sequencing and assembly

The genomes of the four isolates were each assembled into a single large chromosome, between 2.92 and 3.33 Mb. One 39.1 Kb plasmid was recovered for strain AH1, and two plasmids, 23.8 and 39.5 Kb, were recovered from strain GW6. Table 1 contains an overview of the properties of the assembled genomes, which range from 55-58.7 %G+C content and 97-98% protein-coding genes. GW1, GW6, and HF6a each have two *rrn* operons, and AH1 encodes three. A summary of genes associated with sulfur, nitrogen, carbon, and iron metabolisms and genes associated with biofilm formation are presented in Figure 2.

**Figure 2.**
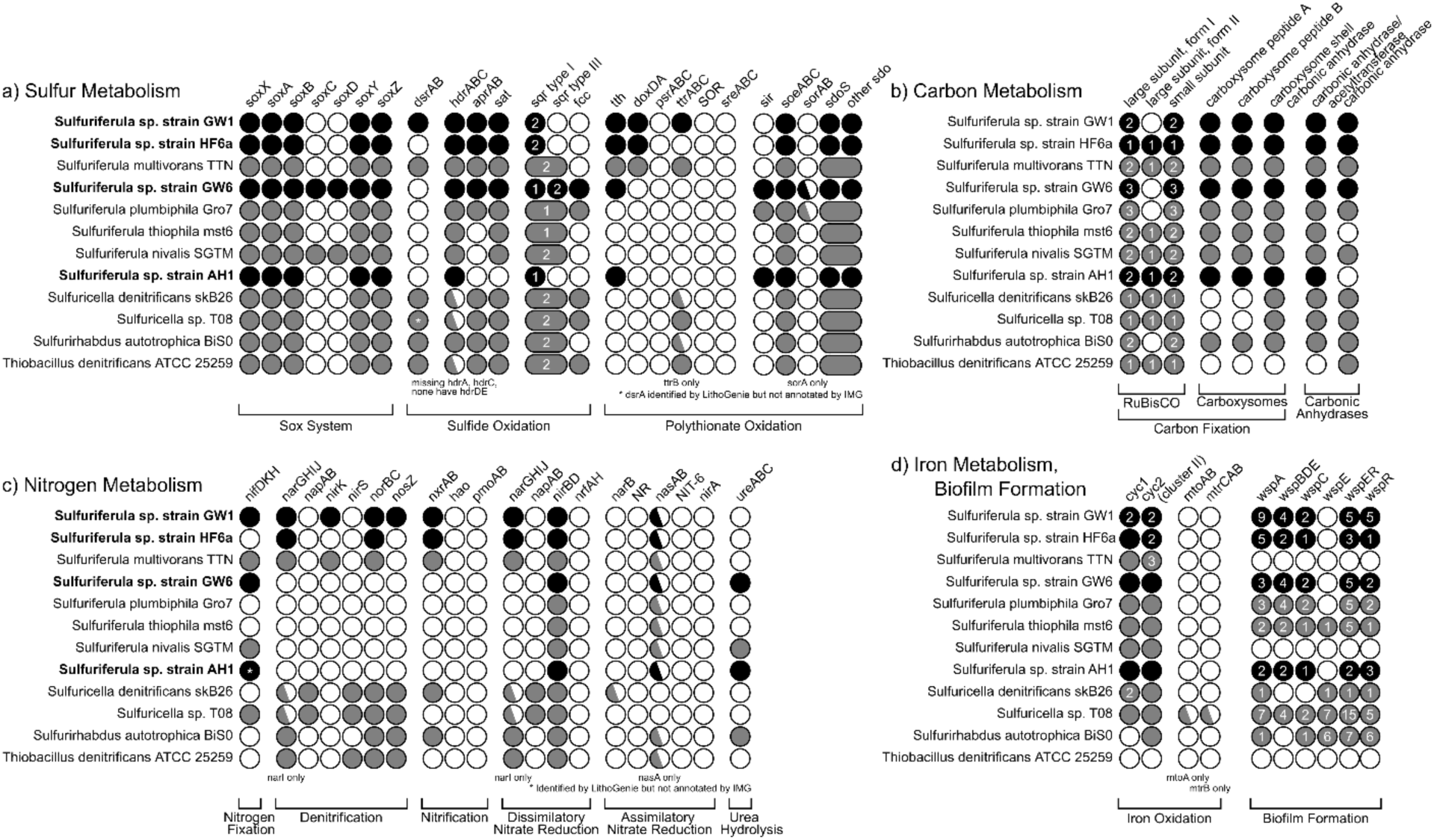
Overview of genes that encode for (a) sulfur, (b) carbon, (c) nitrogen, and (d) iron metabolism and biofilm formation. Filled circles indicate the presence of the gene as annotated by IMG and confirmed by hidden Markov models. A half-filled circle indicates the absence of one or more subunits. The number inside the filled circle indicates the number of copies of the gene found in the genome. Black circles are strains from this study. Larger ovals that overlap the sqr type I/type III and sdoS/other sdo indicate literature sources that did not report those gene clusters as individual genes.

**Table 1.**
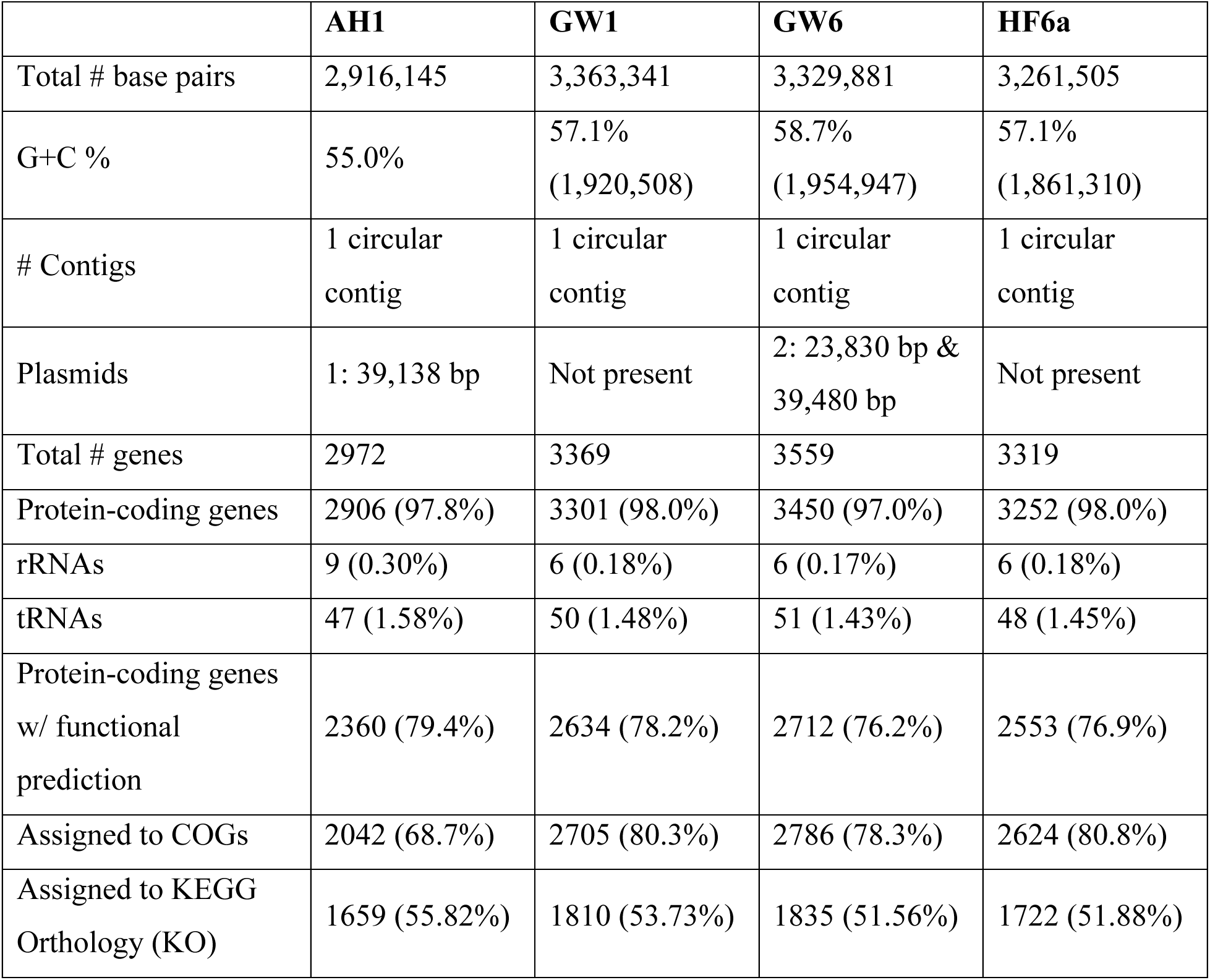
Summary of genome properties.

#### Carbon metabolism

All four strains of *Sulfuriferula* encode complete Calvin cycles for CO_2_ fixation. *S*train AH1 has two copies of RuBisCO form I and one copy of RuBisCO form II, strain GW1 has two copies of RuBisCO form I, strain GW6 has three copies of RuBisCO form I, and strain HF6a has one copy each of RuBisCO form I and II. All four strains have a full complement of carboxysome and carbonic anhydrase-coding genes (Figure 2b). All four strains possess complete glycolysis pathways and incomplete TCA cycles.

#### Nitrogen metabolism

The four strains have diverse capabilities for nitrogen acquisition and dissimilatory nitrogen metabolism (Figure 2c). Strains AH1, GW1, and GW6 encode genes for a molybdenum-dependent nitrogenase typically associated with nitrogen fixation. Strains GW1 and HF6a possess genes *narGHIJ* and *norBC*, indicating that these strains can reduce nitrate and nitrous oxide, respectively. Strain GW1 additionally possesses *nirK* and *nosZ*, providing this strain with a complete set of genes necessary for denitrification. All four strains possess *nirBD* genes that encode dissimilatory nitrate reduction to ammonia. Strains GW1 and HF6a also possess the *nxrAB* genes, indicating the potential for nitrite oxidation. No strains possess complete assimilatory nitrate reduction pathway genes, although all strains possess the *nasA* gene. Additionally, the genomes of strains AH1 and GW6 contain genes *ureABC* that encode urea hydrolysis, indicating an ability to utilize urea as a source of ammonia.

#### Sulfur metabolism

The four strains encode diverse pathways for the oxidation of reduced inorganic sulfur compounds (Figure 2a). All four strains encode sulfide:quinone oxidoreductase (SQR) homologs. SQR is used by all domains of life for sulfide oxidation and/or detoxification. Six structural types of SQR are known; microorganisms encoding at least one representative enzyme from each of type I, III, IV, V, and VI can use sulfide as an electron donor, while type II has not been directly linked to metabolic oxidation of sulfide and may be used for detoxification (43, 44). All four *Sulfuriferula* strains have genes identified as SQR type I, with strains GW1 and HF6a having two copies. Strain GW6 also possesses two copies of the SQR type III, as well as *fcc*, which encodes flavocytochrome-c sulfide dehydrogenase, a structurally-related protein that can also be utilized for sulfide oxidation (45).

Strains AH1, GW1 and HF6a possess partial *sox* systems, with genes *soxAX*, *soxYZ*, and *soxB* but not *soxCD*, while strain GW6 possesses a complete sox system including *soxCD*. In the complete pathway, SoxYZ complexes with thiosulfate to form a SoxY-cysteine S-thiosulfate derivative, which move to sequential reactions with SoxAX and SoxB that oxidize one of the sulfur atoms in the thiosulfate to sulfate, with two electrons transferred to the electron transport chain. SoxCD then oxidizes the remaining sulfane sulfur of the residual SoxY-cysteine-S to sulfate, which is hydrolyzed by SoxB, ultimately generating sulfate and transferring six electrons to the electron transport chain (46). The sox pathway is able to catalyze oxidation of reduced sulfur species besides thiosulfate, including sulfide, elemental sulfur, sulfite, and tetrathionate (47–50), and this ability seems to be facilitated by the modular nature of the reactions (51). While SoxCD is not essential for complete thiosulfate or sulfide oxidation, partial *sox* systems lacking SoxCD are thought to produce elemental sulfur or polysulfides (52). In some organisms with a partial *sox* system, that sulfur is then completely oxidized by the DSR system (53, 54). Only strain GW1 contains genes encoding DsrAB (dissimilatory (bi)sulfate reductase), however, indicating that the other strains either produce S(0) as a final product or use another process to completely oxidize the S(0).

All four strains possess *hdrABC*, which encodes heterodisulfide reductase that is linked to the oxidation of zero-valent sulfur in some bacteria (55). All four strains also possess genes coding for sulfur dioxygenases (*sdo*). Sdo oxidizes the sulfate sulfur in glutathione persulfide (GSSH) to sulfite (56). Homologous genes are found in mitochondria, where mutations on this gene are implicated in a rare hereditary human disease (57). Widely distributed through the Proteobacteria, *sdo* genes are used both in sulfide detoxification and in primary metabolism. The *Sulfuriferula* strains examined here have two types of *sdo* sequences: one set are most similar to the mitochondrial sulfide detoxification ETHE1 genes, while the other set are most similar to the *sdoS* genes found in acidophillic bacteria like *Acidithiobacillus* (Figure 3). Strains AH1, GW1, and HF6a have one copy of each of the two types of *sdo*, while strain GW6 has one copy homologous with ETHE1 and two distinct copies of genes within the *sdoS* group. The strains did not encode other known enzymes involved in S(0) oxidation like sulfur oxygenase reductase (SOR) (58).

**Figure 3.**
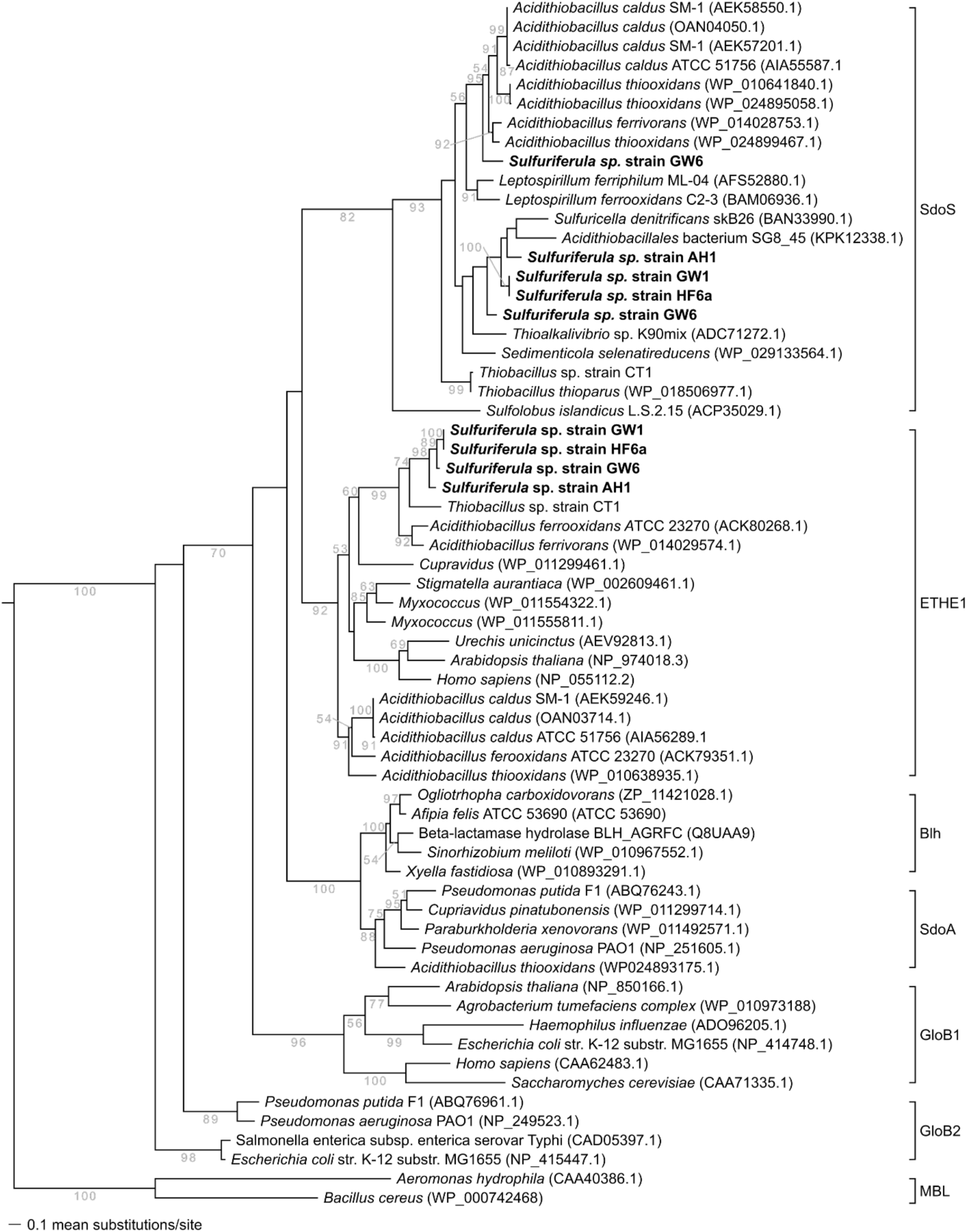
Maximum likelihood analysis of full-length Sdo from isolate organisms, with the genes from the isolates in this study highlighted in bold. The clade of metallo-beta-lactamase (MBL) sequences was used as the outgroup after Liu et al. (56).

All four strains also possess *tth* genes for tetrathionate hydrolase, which catalyzes the oxidation of tetrathionate to thiosulfate, elemental sulfur, and sulfate (59), and *soeABC*, which encodes a sulfite dehydrogenase that allows the transfer of electrons to the quinone pool by the oxidation of sulfite to sulfate (60). Strain GW1 additionally possesses genes *doxDA* for thiosulfate dehydrogenase, which catalyzes the oxidation of thiosulfate to tetrathionate (61), and *ttrABC* for tetrathionate reductase, which catalyzes the oxidation of tetrathionate to thiosulfate (62). Strains AH1 and GW6 possess *sir* genes which code for ferrodoxin:sulfite reductase, catalyzing the assimilatory reduction of sulfite to sulfide (63), and GW6 possesses *sorA*, which encodes the A subunit of sulfide dehydrogenase (64). Strains GW1, GW6 and HF6a possess *sat* genes for sulfate adenylyltransferase and *aprAB* for adenylylsulfate reductase. Both of these proteins typically catalyze the reduction of sulfate (65, 66), but like DsrAB they can run in reverse, oxidizing intermediate organosulfur compounds to sulfate (67, 68).

#### Iron oxidation

All four strains of *Sulfuriferula* described here contain genes that envode for iron-oxidizing cytochromes Cyc1 and Cyc2 (Figure 2d) Cyc1 is a c-type cytochrome likely located in the periplasm and is implicated in iron oxidation in extreme acidophiles (49, 69, 70), while Cyc2 is a candidate iron oxidase found in both neutrophilic and acidophilic iron oxidizers (71, 72). Cyc2 is further divided into three phylogenetically distinct clusters: cluster 1, which has not been experimentally verified but is found in most well-established neutrophilic iron oxidizing organisms; cluster 2, found in *Acidithiobacillus ferrooxidans*, *Ferrovum* spp., *Thiomonas* spp. and other acidophiles, and cluster 3, found in marine iron-oxidizing bacteria and acidophilic *Leptospirillum* spp. (73). The *cyc2* genes in the *Sulfuriferula* strains analyzed here are most similar to those in cluster 2 of the Cyc2 phylogeny of (73).

#### Biofilm formation

All four strains of *Sulfuriferula* contain genes identified as belonging to the wsp system, implicated in biofilm formation (74) (Figure 2d). In *Pseudomonas aeruginosa*, the wsp system senses surface contact and activates genes that promote biofilm formation and regulation (75, 76). Homologs of the *wsp* system are found in the *Beta*- and *Gammaproteobacteria* (77). The four strains are missing *wspE* (Fig. 4d), although studies in *P. aeruginosa* found that mutations on this gene did not affect biofilm formation or morphology (76) although a recent study implicated mutations on *wspE* to the formation of “wrinkly” type biofilms in *Burkholderia* spp. that result in more severe human disease (78).

**Figure 4.**
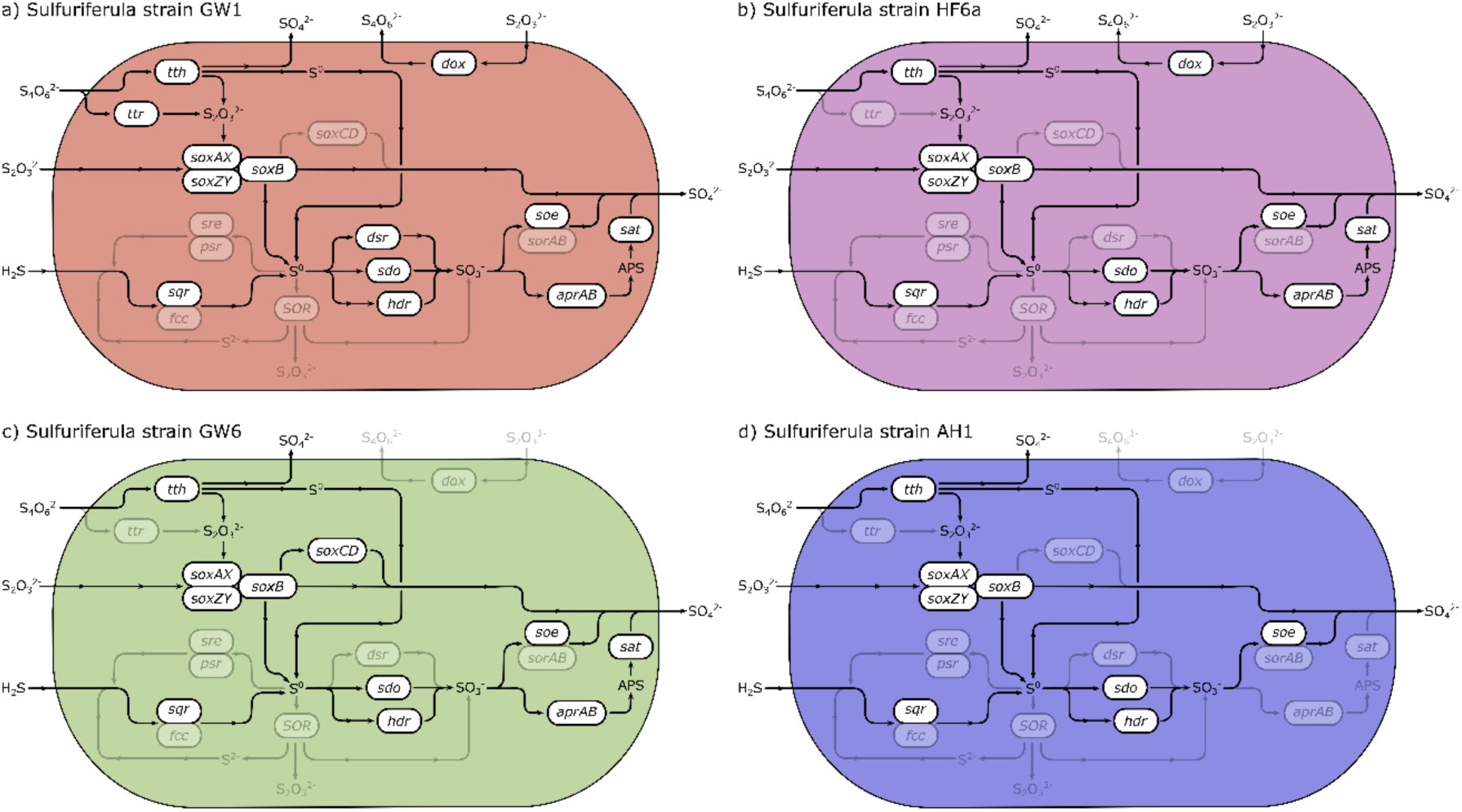
Sulfur oxidation pathways in the four Sulfuriferula strains highlighting differences in metabolic capabilities based on genome reconstruction. Enzymes are as follows: tth, tetrathionate hydrolase (59); dox, thiosulfate oxidoreductase, tetrathionate-forming (61); ttr, tetrathionate reductase (62), sox, multicomponent sulfur oxidation pathway (46); sre, sulfur reductase (83); psr, polysulfide reductase (84); sqr, sulfide:quinone oxidureductase (43); fcc, flavocytochrome c sulfide dehydrogenase (85); SOR, sulfur oxygenase reductase (58); dsr, dissimilatory sulfite reductase (86); sdo, sulfur dioxygenase (56); hdr, heterodisulfide reductase (60); soe, sulfite dehydrogenase (54); sorAB, sulfite dehydrogenase (87); aprAB, adenylylsulfate reductase (88), sat, dissimilatory sulfate adenylyltransferase (89).

## DISCUSSION

### Metabolic and genomic diversity of *Sulfuriferula*

Despite being isolated from the same site and under similar conditions, the four strains of *Sulfuriferula* spp. described here are phylogenetically distinct, with different growth behavior and genomic capabilities for sulfur and nitrogen metabolism. For example, strain AH1, which consistently to the highest density on thiosulfate media under the growth conditions tested here, as well as most rapidly on pyrrhotite (79), also has the fewest genes involved in the oxidation of reduced inorganic sulfur compounds. A schematic overview of the pathway of sulfur through the cell in each strain is shown in Figure 4. In strain AH1, thiosulfate is oxidized to a mixture of sulfate and elemental sulfur by the incomplete sox pathway. Elemental sulfur is thought to accumulate in organisms that lack both *soxCD* and *dsr*, which oxidizes elemental sulfur to sulfate (80, 81). In strain AH1, however, produced elemental sulfur could be oxidized to sulfite (SO_3_^-^) by *sdo* or *hdr*.

Conversely, strain GW6, which is the only strain with the complete sox pathway but otherwise possesses an identical set of sulfur-oxidation genes to AH1, consistently showed the lowest optical density (and therefore presumably the least growth) at all pH conditions tested, suggesting that under these conditions there are other factors affecting growth rate. The addition of organic carbon sources to the growth media (yeast extract, glucose, acetate, succinate) did not increase the growth rate or maximum optical density of strain GW6, so its lower growth rate when compared to other strains is not likely linked to organic carbon or specific nutrient requirements. Strain GW6 does differ from strain AH1 in that it lacks a form II RuBisCO. Since form II is higher efficiency (although lower specificity) than type I (82), the growth rate differences between strains AH1 and GW6 could be a result of differences in efficiency of the carbon fixation pathways.

Strains GW1 and HF6a share many similarities with their closest-related isolate, *S. multivorans* (15), including the presence of *dsrAB* genes (in GW1) and genes for thiosulfate oxidation and reduction (*tth*, *doxDA*). They also share a number of genes encoding proteins involved in nitrogen cycling, including both denitrification and nitrogen fixation abilities. Interestingly, *S. multivorans* lacks the *wsp* genes that are present in all the strains of *Sulfuriferula* described here (Figure 2d), suggesting that biofilm formation is not an important part of its metabolic capabilities. *S. multivorans* was initially enriched by anaerobic thiosulfate oxidation linked to nitrate reduction (15), and so the similarities of the metabolic capabilities of *S. multivorans* and strains GW1 and HF6a suggest that they may also have the ability to grow anaerobically. Although we did not demonstrate anaerobic growth under the conditions tested here, that may be due to specific nutrient requirements that were not present in the media used in this study.

Strain GW6 is most closely related to *S. plumbiphila*, which was isolated on galena (PbS) and was initially found to grow strictly aerobically on H_2_S and H_2_, but not on iron(II), elemental sulfur, thiosulfate, tetrathionate, or other metal sulfides. However, recent work has described growth of *S. plumbiphila* on tetrathionate, thiosulfate, and elemental sulfur (17). Strain GW6 and *S. plumbiphila* both lack most of the genes affiliated with nitrogen cycling, although GW6 possesses genes encoding for a Mo-dependent nitrogenase complex. Strain GW6 is the only isolate described here that possesses the complete Sox system; the only other isolate with genes homologous to *soxCD* is *S. nivalis* (17).

Given the demonstrated growth of other *Sulfuriferula* spp. on elemental sulfur and tetrathionate, we were surprised that all the strains described in this study could not grow on these energy sources. This could be due to specific nutrient requirements not met in the media used here, but it could also reflect specialization in conditions in the mine waste environment, such as growth on acid-soluble sulfide minerals like pyrrhotite or chalcopyrite that are the most abundant sulfides in the Duluth Complex. We were also surprised to find that the strains grew on ferrous iron, given that iron oxidation capability has not been reported for other *Sulfuriferula* strains (15–17, 42). However, it is consistent with the presence of *cyc2* in the genomes of all the isolates. We also note that two of the strains described here, GW1 and HF6a, have the capacity to partially reduce nitrate, with GW1 having a complete set of denitrification genes, showing even more potential metabolic versatility among this genus.

### Implications for the biogeochemistry of Duluth Complex mine waste

All strains in this study grew on the iron-sulfide mineral pyrrhotite as a sole energy source, which is the most abundant sulfide in the weathered rock from which they were isolated. Microbial consortia in biomining or other sulfide mineral-rich environments include organisms that play the role of “oxidant manufacturers”, “acid generators”, and “janitors”, which recycle ferric iron, produce sulfuric acid, and consume low molecular weight organics, respectively (90). All the *Sulfuriferula* spp. characterized here all have the potential to oxidize inorganic sulfur compounds and ferrous iron, indicating that they could either play the role of oxidant manufacturer by generating ferric iron, acid generator by oxidizing intermediate sulfur compounds released during sulfide mineral oxidation, or both, depending on the conditions. Furthermore, pyrrhotite is an acid soluble sulfide that can be oxidized by ferric iron or oxygen, and can also be broken down non-oxidatively by sulfuric acid (91, 92). It is therefore possible that *Sulfuriferula* spp. could directly contribute to the breakdown of pyrrhotite through either its iron or sulfur metabolisms. However, the oxidation of pyrrhotite above pH 4 is not well understood (91, 93–96), so further experimentation is necessary to determine biological pathways of sulfide mineral breakdown in these deposits.

The metabolic diversity of *Sulfuriferula* strains could impact their fitness for, and distribution in, different rock and mineral weathering environments. Previous work found that the *Sulfuriferula* OTUs found in humidity cell experiments on Duluth Complex ore tailings were distinct (<97% 16S sequence similarity) from *Sulfuriferula* OTUs found in sediment from naturally weathered outcrops of ore-bearing Duluth Complex rock (41). In addition to mining-influenced environments (14, 18, 20, 22–24, 53, 97), *Sulfuriferula* spp. have also been found in Antarctic lakes (34), constructed wetlands (28, 29) and hot springs (30, 32, 42), as well as hydrogen sulfide-rich aquatic environments (15–17, 35, 42) The diversity of sulfur oxidation and nitrogen metabolic capabilities in *Sulfuriferula* spp. could result in different strains being more successful in different environments.

For example, humidity cell experiments are typically conducted indoors and have negligible inputs of organic carbon or nitrogen, so *Sulfuriferula* strains with the capability to fix nitrogen (strains AH1, GW1, GW6) will have a competitive advantage over strains that cannot (e.g., strain HF6a). Conversely, in a natural weathering environment, where organic carbon and nitrogen species will be available from plants or soil bacteria, the ability to metabolize oxidized or reduced nitrogen species (like strain GW1, which possesses a complete denitrification pathway) could provide a competitive advantage. Similarly, *S. multivorans* or other organisms which are able to grow heterotrophically or mixotrophically on organic carbon would have an advantage over the obligate chemoautotrophic strains studied here. Although the pH of leachate from experimentally-generated Duluth Complex mine waste remains near-neutral (39, 40), the ability to oxidize iron, particularly if conditions in a waste rock pile or tailings basin became microoxic, could provide an additional source of metabolic energy to these strains.

Strain AH1 displays the most rapid thiosulfate oxidation rates and has also been shown to oxidize sulfide minerals (specifically pyrrhotite, Fe_1-*x*_S, 0 ≤ *x* ≤ 0.125) much more rapidly than other strains (79). This strain appears to have the simplest sulfur oxidation pathway and has the smallest genome out of the four strains examined. This suggests that *Sulfuriferula* sp, strain AH1 is highly specialized for nutrient-poor sulfide mineral weathering environments, and indicate that strain AH1 may have applicability in biomining and mine remediation. *Sulfuriferula* spp. are sensitive to organic acids, and do not appear to be able to utilize organic carbon, and so their activity in tailings or other mine waste environments could be controlled by the application or removal of organic carbon compounds. If the desired goal is to prevent sulfide mineral oxidation, adding organic compounds to the tailings pile would likely result in the chemolithoautotrophic *Sulfuriferula* spp. being outcompeted by heterotrophic organisms (e.g., 41, 98–100). Conversely, if the goal is to accelerate sulfide mineral dissolution, *Sulfuriferula* spp. could be enriched by maintaining an environment depleted of organic carbon. Further experimentation will be necessary to determine the biotechnological potential and explore specific applications for this group of bacteria across the mining lifecycle.

## METHODS

### Isolation and identification of *Sulfuriferula* strains

Isolates AH1, GW1, and HF6a were isolated on solid thiosulfate media containing 10 g L^-1^ NaS_2_O_3_·5H_2_O, 1.5 g L^-1^ KH_2_PO_4_, 4 g L^-1^ Na_2_HPO_4_, 0.4 g L^-1^ (NH_4_)_2_SO_4_, 0.4 g L^-1^ MgSO_4_·7H_2_O, 0.04 g L^-1^CaCl_2_·2H_2_O, trace element solution (101), and 1.5% agar. Strain GW6 was isolated similarly, but with urea provided as the sole N source.

### Growth conditions

Growth of *Sulfuriferula* strains was assessed on liquid thiosulfate media, either buffered with phosphate (containing 10 g L^-1^ NaS_2_O_3_·5H_2_O, 1.5 g L^-1^ KH_2_PO_4_, 4 g L^-1^ Na_2_HPO_4_, 0.4 g L^-1^ (NH_4_)_2_SO_4_, 0.4 g L^-1^MgSO_4_·7H_2_O, 0.04 g L^-1^CaCl_2_·2H_2_O, and a trace element solution (101)) or MES (reducing KH_2_PO_4_ and Na_2_HPO_4_ to 16 and 6 mg L^-1^, respectively, and adding 20mM MES buffer) to the appropriate pH. Growth was assessed qualitatively by measuring optical density (OD) at 600nm and 450nm. Potential utilization of different carbon sources was tested by growing the isolates on LB broth, 1:10 LB broth, or the phosphate-buffered thiosulfate media with 1% yeast extract, 5mM glucose, 5mM acetate, or 5mM succinate. Growth experiments were conducted in clear borosilicate test tubes or serum bottles, and growth at starting pH values of 4, 4.5, 6, 7, 7.5, and 8 was compared using MES buffered thiosulfate media. We did not test growth below pH 4 because thiosulfate hydrolyzes abiotically and we could not identify an alternative growth substrate, and because of precipitate in the media above pH 8. We do not report specific growth rates because optical density measurements are impacted by the formation of S^0^ during hydrolysis of thiosulfate in the media, and by the formation of other precipitates. To track pH and sulfate production during growth, pH was measured using a LAQUAtwin pH-22 handheld pH meter (HORIBA, Kyoto, Japan) and sulfate was measured in duplicate by barium sulfate (BaSO_4_) precipitation and absorbance at 450nm.

Growth on hydrogen sulfide and ferrous iron was evaluated with a gradient tube culturing technique (102–104). Gradient tubes were prepared using the phosphate buffered media base as described above (1.5 g L^-1^ KH_2_PO_4_, 4 g L^-1^ Na_2_HPO_4_, 0.4 g L^-1^ (NH_4_)_2_SO_4_, 0.4 g L^-1^MgSO_4_·7H_2_O, 0.04 g L^-1^CaCl_2_·2H_2_O, and trace elements). The plug was prepared with media based, 1% agar, 2 mM NaHCO_3_, and either 2 mM NaS_2_ or 10 mM FeSO_4_. The overlay was prepared with media base and 0.25% agar. Tubes were inoculated after allowing the gradients to develop for 5 days, and growth was evaluated based on the presence of a growth band as well as cell visualization using DAPI (Supplementary Figure S2). We found that it was necessary to add HCO_3_^-^ to ensure cell growth, as well as high Fe^2+^ concentration to ensure effective gradient development in the iron gradient tubes. Growth on pyrrhotite was evaluated by adding 0.5 g of crushed pyrrhotite, prepared as in (79, 105), to thiosulfate-free growth media with both phosphate and MES as buffers.

### Genome sequencing, assembly, and annotation

Cells of the strains investigated in this manuscript were collected from batch cultures onto 0.2μm-pore filters and genomic DNA (gDNA) was isolated from the filters using the PowerSoil DNA isolation kit (Qiagen). For strain AH1, SMRTbell libraries were prepared with 20μg of high molecular weight DNA, and sequenced on a Pacific Biosciences (PacBio) RSII platform. For other strains, SMRTbell libraries were prepared with 2-3 ug of high molecular weight DNA was prepared with a 10 Kb insert size at the University of Minnesota Genomics Center (UMGC), and pooled and sequenced on a PacBio Sequel platform. Assembly of AH1 was described in (106), and the other genomes were assembled using HGAP version 6 with default parameters.

Sequences were uploaded to IMG/MER (107) and annotated using the JGI Integrated Microbial Genomes (IMG) Pipeline (108). Presence and function of genes involved in carbon and nitrogen metabolic pathways were determined using KEGG Ortholog annotation (109). Genes involved in sulfur metabolism, iron metabolism, and biofilm formation were initially determined using KEGG Orthology using GhostKOALA (109) and were confirmed using hidden Markov models (HMM) with HMMer v. 3 (110). HMMs for genes in the *SoxB*, *SoxC*, and *SoxD*, as well as *Sqr*/*Fcc* pathways were constructed by aligning a database of sequences using TCOFFEE’s Expresso aligner (111) and using HMMER to create the model out of the aligned sequences. Databases for *soxB* were generated using sequence compilations from (112, 113), databases for sqr were generated from (6, 43), and databases for sdo were generated from (56, 114). The isolate genomes were then converted from DNA to proteins using Prodigal v. 2.6.3 (115) and screened for matches using hmmsearch with an E-value cutoff of 10^-7^. All other HMMs used were part of the MagicCave package (116, 117).

### Phylogenetic analysis

For phylogenetic analysis of 16S rRNA genes, sequences were aligned in ARB (118) using the Silva database v. 132 (119). Sequences were filtered so that all *Sulfuriferula* sequences were the same length and positions with more than 50% gaps removed (final alignment 1449 positions). Neighbor joining analyses were performed with the ARB-implementation of phylip (120) with Jukes-Cantor-corrected distance. SDO phylogenies were created with maximum likelihood analysis using RAxML v.8.0.24 (121) with the LG model of amino acid substitution (122) with observed amino acid frequencies and the fraction of invariant sites and the shape parameter (α) value estimated from the data. Phylogenetic analysis of SDO genes was performed with representative sequences from (56, 114), and using clade names from those publications.

### Accession numbers

The genomes of the *Sulfuriferula* strains in this study are available at the National Center for Biotechnology under Bioproject accession PRJNA1160330. The specific accessions for each strain and associated plasmids are CP021138-CP021139 (str. AH1), CP170197 (str. GW1), CP170194-CP170196 (str. GW6), and CP170193 (str. HF6a).

## ACKNOWLEDGEMENTS

We thank the Minnesota Department of Natural Resources Division of Lands and Minerals for facilitating sampling of the weathered rock samples from which the strains were isolated, especially Z. Wenz, K. Lapakko, M. Olson, S. Koski and P. Geiselman for access and assistance with sample collection. Special thanks to A. Baloun and A. Hua for assistance with culturing, and to B. Auch and the staff at the UMGC for assistance with genome sequencing. This work was supported by University of Minnesota MnDRIVE initiative and the University of Minnesota BioTechnology Institute. This work was supported by University of Minnesota MnDRIVE initiative and the University of Minnesota BioTechnology Institute.

## Notes

### Competing Interest Statement

The authors have declared no competing interest.

### Summary of Updates

Add supplementary materials and update author information

