## Supplementary Figures for "*Sulfuriferula* spp. from sulfide mineral weathering environments have diverse sulfur- and iron-cycling capabilities"

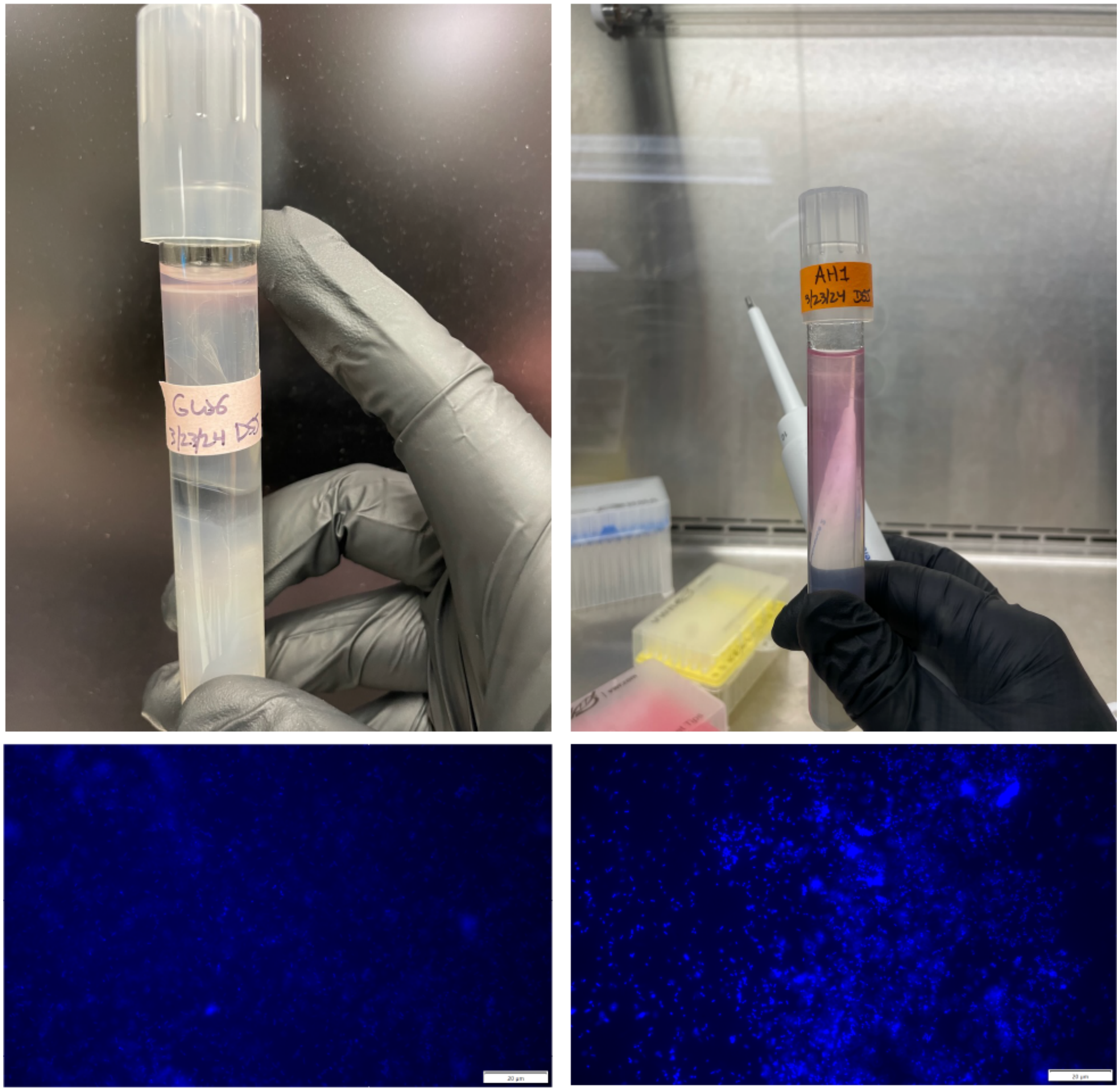


**Supplementary Figure S1.** Representative photos of gradient tubes showing growth on $H_2S$ (upper left) and $Fe^{2+}$ (upper right). Growth was verified by DAPI staining cells from the growth bands at the top of the cultures (lower two images).

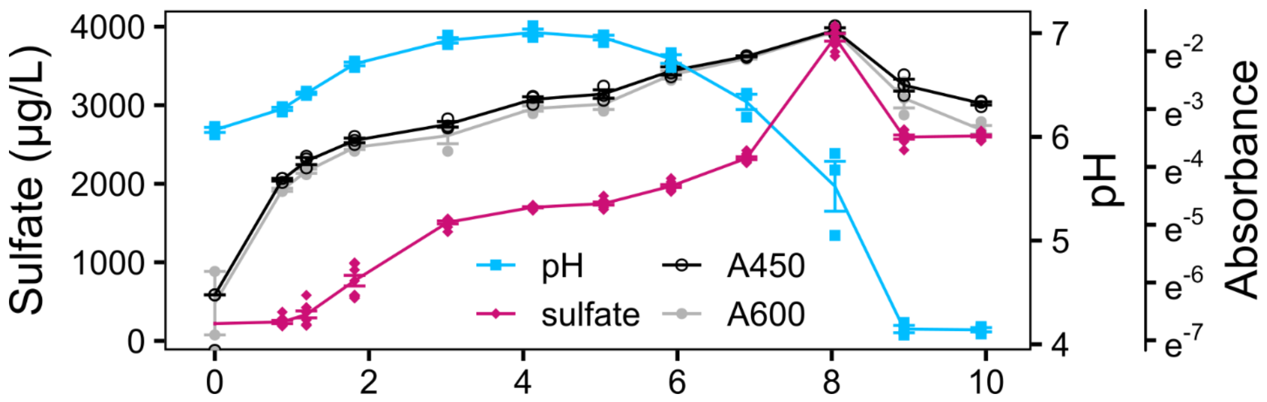


**Supplementary Figure S2.** pH, aqueous sulfate concentration, and absorbance at 450nm (A450) and 600nm (A600) for *Sulfuriferula* sp. strain AH1 grown on thiosulfate in pH 6.0 growth media.
